# Active smelling in the American cockroach

**DOI:** 10.1101/2022.11.23.517676

**Authors:** Antoine Hoffmann, Einat Couzin-Fuchs

## Abstract

Motion plays an essential role in sensory acquisition. From changing the position in which information can be acquired to fine scale probing and active sensing, animals actively control the way they interact with the environment. In olfaction, movement impacts the time and location of odour sampling as well as the flow of odour molecules around the olfactory organs. Employing a high-resolution analysis, we investigate how the insect antennae interact with the olfactory environment in a species with a well studied olfactory system – the American cockroach. Cockroaches were tested in a wind-tunnel setup during the presentation of odours with different attractivity levels: colony extract, butanol and linalool. Our analysis revealed significant changes in antennal kinematics when odours are presented, including a shift towards the stream position, an increase in vertical movement and high-frequency local oscillations. Nevertheless, the antennal shifting occurred predominantly in a single antenna while the overall range covered by both antennae was maintained throughout. These findings hold true for both static and moving stimuli and are more pronounced for attractive odours. Furthermore, we find that upon odour encounter, there is an increased occurrence of high-frequency antennal sweeps and vertical strokes, which are shown to impact the olfactory environment’s statistics directly. Our study lays out a tractable system for exploring the tight coupling between sensing and movement, in which antennal sweeps, in parallel to mammalian sniffing, are actively involved in facilitating odour capture and transport, generating odour intermittency in environments with low air movement where cockroaches dwell.

## 3 Introduction

Olfaction is crucial in almost every aspect of life: finding food and assessing its quality, avoiding danger, communication, and mating. Odour molecules are transported from their sources by currents of air (or water) that vary in flow and turbulence. Both the physical properties of the airflow (or other carrying media) and the chemical attributes of the odor, such as volatility (Cometto-Muñiz et al. (2003)), impact its dispersal, often creating complex spatio-temporal concentration distributions (Moore and Crimaldi (2004); Murlis et al. (2000); Pannunzi and Nowotny (2019)). Nevertheless, despite the structural complexity of natural odour landscapes, animals successfully utilise information from these concentration distributions to track odour plumes (Murlis et al. (1992); Vickers (2006)), localise their sources and segregate concurrent odours that arise from different objects (Baker et al. (1998); Sehdev and Szyszka (2019)).

An important aspect of olfactory behaviour, as commonly acknowledged in the recent literature, is the active nature of interactions between the sensing body and the olfactory environment (Crimaldi et al. (2021); Wachowiak (2011)). Examples for such interactions are exhibited in a variety of distinctive odour-guided behaviours such as casting across plume edges (Kennedy (1983); WILLIS and BAKER (1984)), sniffing (Wachowiak (2011)), antennal movements (antennule “flicking” in crustaceans Koehl et al. (2001), antennal movements in insects Claverie et al. (2021); Huston et al. (2015); Lei et al. (2022)) and wing fanning (Li et al. (2018)). A full mechanistic understanding of these interactions and their role in olfactory processing is still in early stages however, especially when we compare it with the highly detailed knowledge on the olfactory circuits themselves (Galizia and Rössler (2010)). To this end, we note the importance of tractable model systems in which the bidirectional interactions between motion and sensing could be investigated in detail. In this study, utilising an insect preparation with a well-described olfactory system and highly mobile antennae, the American cockroach, *Periplaneta americana*, we study how the olfactory environment impacts, and are impacted by, antennal movements.

Cockroaches are mainly ground-dwelling insects that favour humid, confined spaces in low light conditions. Their exceptionally long and mobile antennae, covered with olfactory sensilla distributed along the entire length, provide a wide working range for odour detection (Lockey and Willis (2015)). Odour information is transmitted into the antennal lobe where information about odour identity and quantity can be evaluated based on the population of activated glomeruli (Watanabe et al. (2010)). Less is known about how the spatial structure of the odour environment is encoded. Presumably, both bilateral comparisons of antennal inputs (“tropotaxis”, Borst and Heisenberg (1982); Duistermars et al. (2009)) and temporal evaluation (“klinotaxis”, Lockey and Willis (2015); Willis and Avondet (2005)) are involved. An antennotopic neural arrangement that allows spatial mapping along the insect antenna, even without movement, has so far only been shown for the specialised pheromone-processing macroglomerulus pathway (Nishino et al. (2018); Paoli et al. (2020)). It is unknown, however, whether cockroaches are able to behaviourally utilise this neural organisation for navigation, and/or if similar antennotopic organisations exist for other odorants as well (Nishino et al. (2018); Paoli et al. (2020)).

Spatial evaluation and successful source localisation of all odours, nevertheless, rely on successive behavioural sampling. These typically follow either a reactive method - in which odour encounters elicit a switch in behaviour (e.g from crosswind casting to up-wing surging, Cardé and Willis (2008)), and/or a strategic one - where odour information is integrated over time (Pang et al. (2018); Voges et al. (2014)). When navigating toward distant odour sources cockroaches utilise wind direction information and exhibit casting movements across the lateral boundaries of wind-borne plumes (Willis and Avondet (2005); Willis et al. (2008)) in a similar manner to odour-source tracking in flying insects (Talley et al. (2022)). Motivated to further understand how, at the local scale, the antennae actively participate in sampling and mapping the olfactory space, we study antennal kinematics during static and moving odour encounters.

This paper is structured as follows: we start with the characterisation of spatial antennal sampling to investigate whether and how antennal movements are modified with respect to the location and nature of surrounding odorants. Our working hypothesis is that, as in other types of search behaviours (Mehlhorn et al. (2015)), exploration-exploitation trade-offs also hold for antennal movement when scanning an odour space. We therefore expect efficient scanning strategies to include both episodes of localised movement, focused on a position of interest, and also wide range scanning of the rest of the available space. We then continue to characterise the fine-scale temporal dynamics of antennal oscillations and discuss the relevance of the observed odour-driven changes for odor transduction and intermittent sampling. The final part of our paper focuses on the impact antennal movements have on the olfactory environment itself.

## 4 Material and Methods

### 4.1 Experimental Setup

Adult males of the American cockroach, *Periplaneta americana*, were kept on a 12/12 h light/dark cycle at 24°C and 65% humidity in the animal facility of the University of Konstanz (Germany). Male cockroaches were collected and briefly immobilised through cold treatment to facilitate handling. Animals were tethered on a glass surface in a 20 × 20 × 100 cm Plexiglas wind tunnel (custom-design, Fig. 1A), using a thin “leash” of a plastic tube wrapped around the thorax and attached to a metal rod for fixation. Care was taken to place the animal in a natural position that allows flexible stepping on a glass surface rendered slippery with a thin layer of glycerol. Behavioural experiments started after 20 min of acclimatization. We performed three experimental protocols for a given animal: A static frontal odour stimulus centred on the animal’s head (“position 1”), a static frontal stimulus on the side, 3 cm from the midline, which is well within antennal reach (“position 2”), and a moving frontal stimulus shifting from position 1, via position 2, to the very edge of the antennal reach (“position 3”). Three odours were delivered in a pseudo-randomised order (colony odour, butanol, linalool).

**Figure 1.**
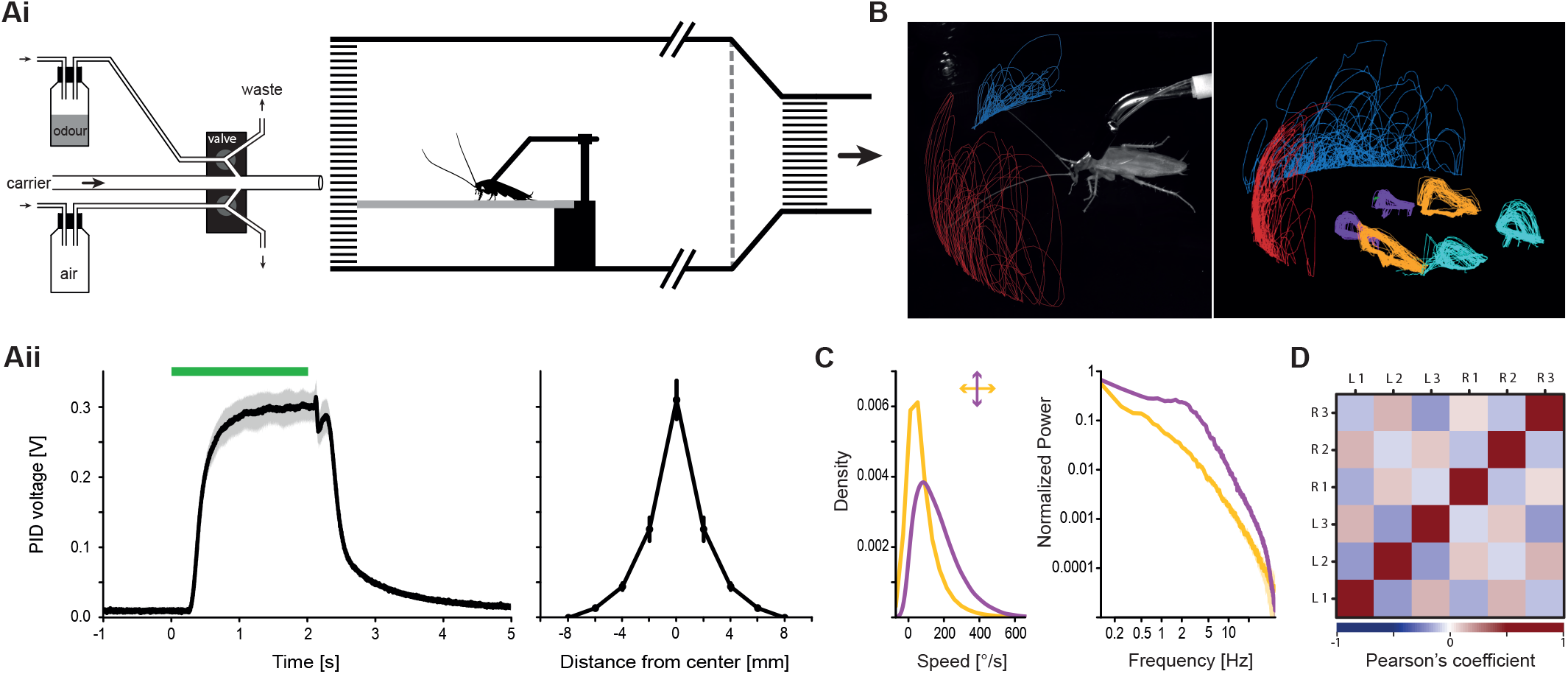
Wind tunnel setup to test active cockroach olfaction. (Ai) Schematics of the odour delivery device Raiser et al. (2017) and wind tunnel. (Aii) Odour stimulus. Left: Photo-ionisation detector (PID) signal as a proxy for the amount of odour molecules for a 2 second pulse (green bar). The delay between the valve trigger and the onset of the odour was compensated for in every subsequent analysis. Right: PID signal for the width of the odour pulse. Mean ± s.e.m.; N = 5. (B) Left: View from one of the cameras from the 3D tracking system of a male cockroach in the wind-tunnel. The animal was free to move its head and limbs on a slippery glass surface. Right: 3D tracking example for the antennae tips, head, and leg movements. (C) Overall analysis of antennal kinematics from pooled trials shows movement to be generally slower in azimuth (yellow) and faster in elevation (purple). Plotted are the distributions of angular antennal speeds with the corresponding Fourier spectra (N = 100). (D) Pairwise correlation matrix of leg kinematics, pooled from all trials, show that leg movement generally follow a typical cockroach-walking tripod gait. R1-R3 represent front to hind right legs, L1-L3 left legs (Pearson’s correlation coefficients; N=330).

**The wind tunnel** was designed to create a laminar air flow for delivering spatially and temporally precise odour stimuli. It includes a rotary fan placed in the exhaust tubing to create an air suction which is evacuated to the laboratory ventilation system. To reduce turbulence caused by the fan, a 2cm thick aluminium honeycomb mesh and a net was placed at the end of the tunnel. An additional honeycomb at the entrance of the tunnel assured a laminar air flow in the experimental recording area (see Fig. 1A). The air speed was set at 0.15-0.2 m/s for all behavioural experiments, far below the typical running speed of the American cockroach.

### 4.2 Odours

Odours were delivered via a custom-made olfactometer based on Raiser et al. (2017) which allows stable, flow-compensated delivery of odorants for extended periods of time (fig. 1B). Three odours were presented to the animals: butanol (Sigma-Aldrich, product code: 19422), a compound found in rotting food as a product of fermentation processes, linalool (Sigma-Aldrich, product code: L2602), a floral compound, and a “colony extract” collected from the substrate in the rearing colonies which is a mixture of faeces odorants, pheromones, and likely other odorants from conspecifics. Odour dilutions (10-2 for butanol and 10-1 linalool) in mineral oil (Acros Organics, product code: 124020010) were chosen due to their overall similar strength of response in the cockroach antennal lobe (Paoli et al. (2020) and unpublished data from the lab). 200 μL of butanol and linalool were placed in 20 mL vials (Schmidlin Labor + Service GmbH, product code: 5520090873) sealed with Teflon septa (Schmidlin Labor + Service GmbH, product code: 5520030142) to be used for delivery and all odorants were regularly renewed to avoid depletion. The stimulus sequences were pre-programmed for accurate repeatability in custom LabVIEW software (NI, Austin, TX, USA) which controls a compact RIO system equipped with a digital I/O module NI-9403 (National Instruments, Austin, TX, USA). For experiments with a moving stimulus, the outlet of the delivery device was shifted laterally on a rail via an Arduino-actuated stepper motor and a custom-written script, synchronised with the odour stimulus sequence. The profile of odour distribution in the wind-tunnel was tested with a Photo-ionisation Detector at its lowest pump setting (Mini-PID model 200A - Aurora Scientific Inc., Ontario, Canada; Recordings with the Spike2 software, Cambridge Electronic Design Limited, Cambridge, England) to ensure a sharp onset and offset in time and space (sharp rise and decay times with a stable plateau within, and a narrow stimulus width, Fig. 1Aii).

### 4.3 Recording & tracking

The experimental apparatus was placed in full darkness, illuminated only with infrared lights (850nm wavelength, model IN-907, Instar, Germany) to avoid visual cues. Recordings were made via the Motif system developed by LoopBio (Loopbio GmbH, Vienna, Austria; http://loopbio.com/recording/): Three calibrated and synchronised Basler cameras (Model acA1920-150um, Basler AG, Germany; Lenses by Kowa Optimed GmbH, Germany) equipped with IR long pass filters were placed to capture the animal from three different views at 100 frames per second. Uniform lighting conditions for the video recordings were obtained by arranging infrared lights (850 nm wavelength) around the recording area. Tracking and 3D reconstruction of the animal’s legs, head and antennae tips was achieved with the LoopBio software “Loopy” (http://loopbio.com/loopy/). The tracking data was then further processed with custom-written scripts in R (R version 4.2.0., https://cran.r-project.org/). All metrics and analyses were calculated from these 3D coordinates in R.

### 4.4 Behavioural data Analysis

**Antennal angles** were computed by approximating each antenna as a straight line from the tracked head point to the tracked antennal tip (cockroaches cannot voluntarily bend their antennae). In the horizontal plane (azimuth), the angle between these lines and the animal’s midline (a virtual line through the animal’s tracked head point) was calculated, such that 0° corresponds to an antenna exactly parallel to the midline. Negative angles are left and positive angles right. In the vertical plane (elevation), the angle between that same virtual horizontal midline and the approximated antenna was calculated such that negative values correspond to the tip being below, and positive values above, the head line.

**Antennal coordinates** were normalised to compare antennal positions across trials and animals. We centred the X and Y coordinates (left-right and back-front, respectively) around the head point such that the coordinates (X,Y) = (0,0) corresponds to the head position on the horizontal plane. The Z coordinates (vertical axis) were shifted such that zero corresponds to head level. For the analyses in Fig. 3 and Fig. 4, we then normalized the coordinates to their respective maximum absolute values within a trial, thus effectively normalising by the antennal length of each animal. Metrics in these analyses were calculated within this new coordinate system, and thus expressed as a proportion of the antennal length (antennal lengths of our test animals approximately ranged from to 45 to 55 mm). For the spatial ‘heatmaps’ of tip position distributions in Fig. 2, the coordinates were normalised by the maximum possible coordinate across trials in the dataset.

**Figure 2.**
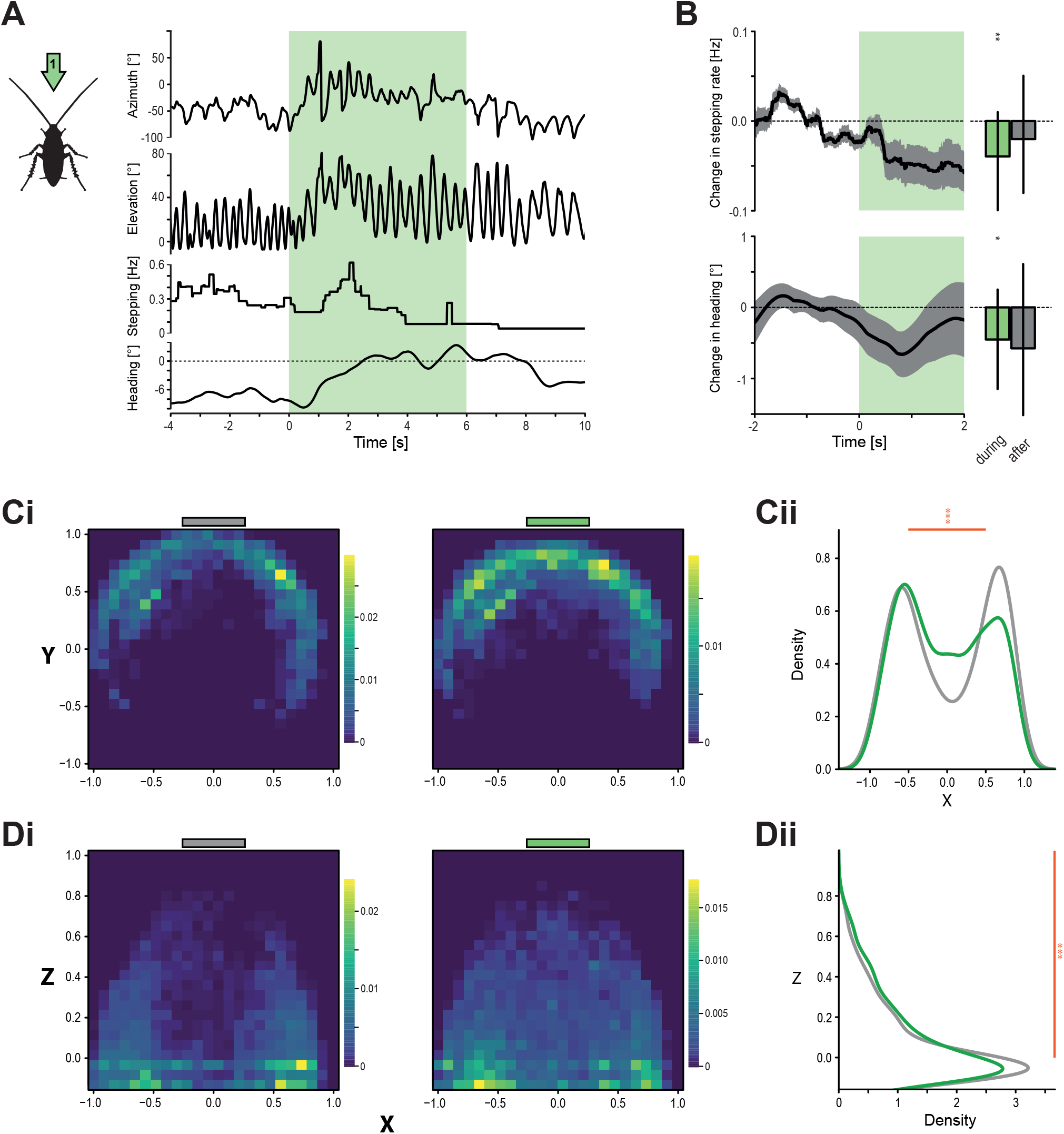
General odour-induced behavioural responses. (A) Left: Schematic view of the stimulus in position 1 (centre). Right: Single trial example with time-series for antennal angles in azimuth and elevation, mean stepping rate, and heading angle (smoothed; negative and positive values indicate left turn and right turn respectively). The shaded green time window indicates the odour stimulus. (B) Top: Change in stepping rate with colony odour, compared to the pre-stimulus air (n=40 trials, N=29 animals). Bottom: Change in absolute heading, compared with the pre-stimulus heading (negative values indicate that the animals are moving their heads towards the stimulus; n=47 trials, N=30 animals; mean time-series ± s.e.m. and model means with credible intervals). (C) Horizontal antennal tip distributions before (left, grey bar) and during (right, green bar) a colony odour stimulus (i) Presence density maps in the normalised coordinate system (zero = head position, ±1 = maximum coordinate across trials). Colours indicate relative proportions of observations per pixel. Grey and green bars above indicate the location of the air/odour stimulus. (ii) Antennal tip distributions on the X axis (left-to-right) before and during the odour stimulus (grey and green curves respectively). (D) Vertical antennal tip distribution before (left, grey bar) and during (right, green bar) colony odour stimuli. (i) Presence density maps in the normalised coordinate system (zero = head position, +1 = maximum coordinate across trials). Colours indicate relative proportions of observations per pixel. Antennal tip distributions on the Z axis (ground-to-top) before and during the odour stimulus. In C and D: n=44 trials, N=24 animals, and asterisks indicate the certainty levels of the odour means to be different from the air control (*: ≥ 90%, **: ≥ 95%, ***: ≥ 99%)

**The distance** to the stream (Fig. 3 & 4) was calculated using the absolute Euclidean distance between the antennal tip and the centre line of the stimulus stream. **The antennal range** (Fig. 3 & 4) corresponds to the absolute maximum difference between horizontal positions of each individual antenna. **The overall range** corresponds to the absolute maximum difference between the left-most and the right-most positions of both antennae combined. **Pearson’s correlation** coefficients (Fig.S 2) between the “centre of sweeps” of each antenna and the centre of the stream were additionally calculated. The centre of sweep corresponds to the average X coordinate at a given time point, calculated with a moving average.

**Figure 3.**
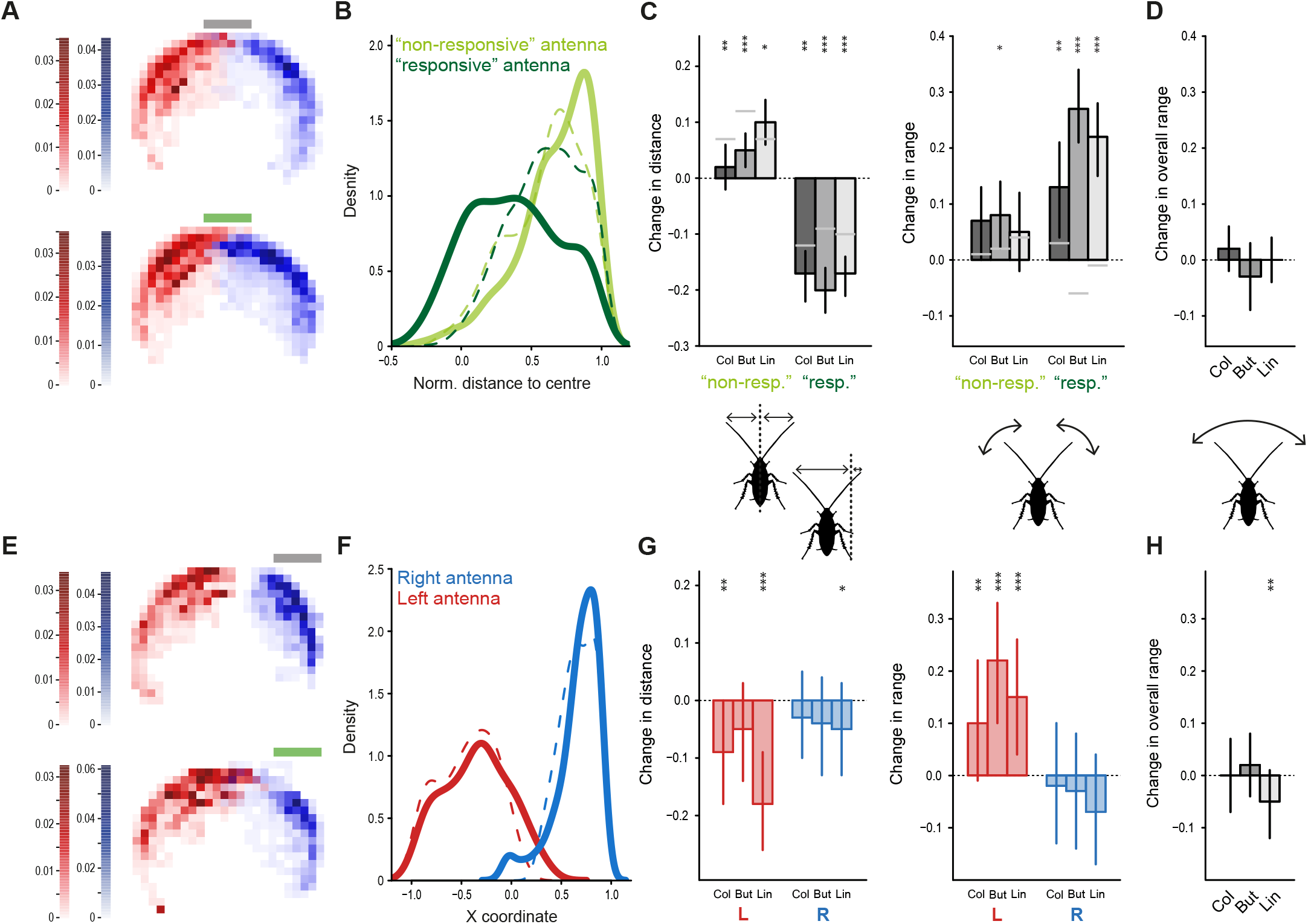
Trade-off between local sampling and global scanning and asymmetries in antennal movement. (A) Horizontal antennal tip distributions before (top) and during (bottom) a colony stimulus, delivered centrally in position 1 (red & blue = left & right antennae). Same data as in Figure 2C. (B) Distributions of the horizontal distances between the antennal tip and the stimulus centre. In each trial, the antennae were labelled as “responsive” and “non-responsive”, based on their movement towards the odour (see Methods). Dashed lines correspond to the air control, full lines correspond to the distributions during the odour stimulus (here, colony). (C) Left: Odour-induced changes in distance to position 1 for each antenna (“responsive” and “non-responsive”) and each odour (means and credible intervals). Right: Odour-induced changes in antennal range for each antenna (“responsive” and “non-responsive” identities transferred from the left panel) and each odour (model means and credible intervals). Grey lines indicate baseline control changes (see Methods). Asterisks indicate the certainty levels of each mean to be different from its respective control (*: ≥ 90%, ≥ **: ≥ 95%, ***: 99%). (D) Changes in overall antennal range for the three tested odours (means and credible intervals; n=44 trials, N=24 animals for Col; n=46, N=25 for But; n=48, N=25 for Lin). (E) Same as in (A), but for an odour stimulus in position 2 (side stimulus). (F) Distributions of left (red) and right (blue) antennal tips during pre-stimulus air (dashed lines) and stimulus (colony, full lines). (G) Left: Odour-induced changes in distance to position 2 for each antenna and each odour (model means and credible intervals). Right: Odour-induced changes in antennal range for each antenna and each odour (model means and credible intervals). (H) Odour-induced changes in overall antennal range for each odour (model means and credible intervals). In G and H, asterisks indicate the certainty levels of each mean to be different from zero (*: ≥ 90%, **: ≥ 95%, ***: ≥ 99%) with n=20 trials, N=11 animals for Col and But; n=21, N=11 for Lin.

**Figure 4.**
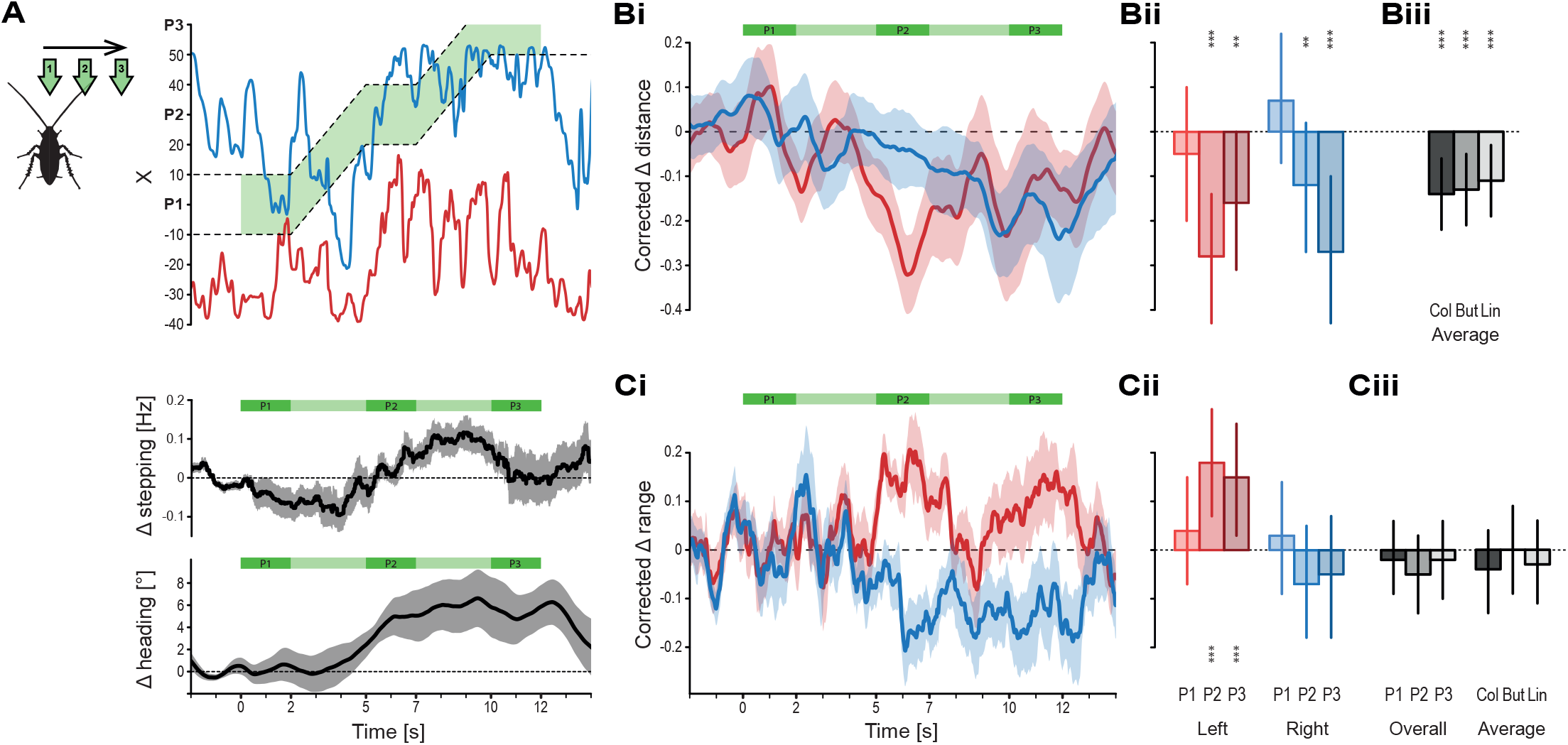
Stream following behaviour. (A) Left: Schematic view of the stimulus shift with three static periods in position 1 (P1 - centre), position 2 (P2 - side) and 3 (P3 - extreme side). Top right: Example trial of antennal tip positions on the X axis (left to right) over time. Dashed lines delineate the location of the stream. Green shading represents the time of the odour stimulus (here, colony). Middle: Change in stepping rate over time (mean ± s.e.m.). Bottom: Change in absolute heading direction (left-to-right) over time (mean ± s.e.m.). (B) Changes in distance of antennal tip positions to the stimulus centre. Negative values indicate larger displacements toward the odour compared with the air control. (i) Time-series of control-corrected changes in distance to the moving stream centre for both antennae for the colony odour (ii) Model means and credible intervals of control-corrected changes in distance for each antenna in the static positions P1-3 (colony). (iii) Overall model means across P1-P3 with credible intervals for each odour. (C) Changes in antennal range. Positive values indicate larger ranges than for the air control (i) Time-series of control-corrected changes in range for both antennae for colony (ii) Model means and credible intervals of control-corrected changes in range for each antenna in the static positions P1-3 (colony odour). (iii) Left: Model means of overall range (total range of the two antennae combined) and credible intervals for the static positions for colony. Right: Overall model means across P1-3 and credible intervals for each odour. N = 13 for each odour. In B and C, asterisks indicate the certainty levels of each mean to be different from zero (*: ≥ 90%, **: ≥ 95%, ***: ≥ 99%).

**Spatial heat maps** of antennal tips were generated from the centered and normalised coordinates. The (X,Y,Z) = (0,0,0) position in each heat map thus corresponds to the head position (Fig. 2). The sum of observations of antennae tip positions across trials was computed in 24 bins for each axis. The heat maps were then individually normalised by their overall observation sums for comparison. Density curves correspond to the empirical kernel probability density function (*density* function from the base *stats* package in R, the same bandwidth parameter value was used for the curves of a given panel). The area under each curve thus equals 1.

#### Frequency analysis

The average frequency spectra in Fig. 1 are based on a Fast Fourier Transform (*spec* function, *seewave* package in R (Sueur et al. (2008))). The spectral analysis in fig. 5 is based on a continuous wavelet analysis (*analyze*.*wavelet* function, *WaveletComp* package in R (Roesch and Schmidbauer (2018))). The continuous wavelet power spectra of angular time-series were averaged in 1s time-bins and pre-defined frequency bands ([0.5-1), [1-2), [2-3), [3-5), [5-10) Hz). The power values for each frequency band were then scaled as follows: Δ*P/P* = (*P*_*t*_ − *P*_0_)*/P*_0_, where *P*_*t*_ is the power of a given time-bin and *P*_0_ is the average power of a given frequency band in the pre-stimulus window. The resulting metric thus quantifies changes in wavelet power for each frequency band as a proportion of the pre-stimulus average.

**Figure 5.**
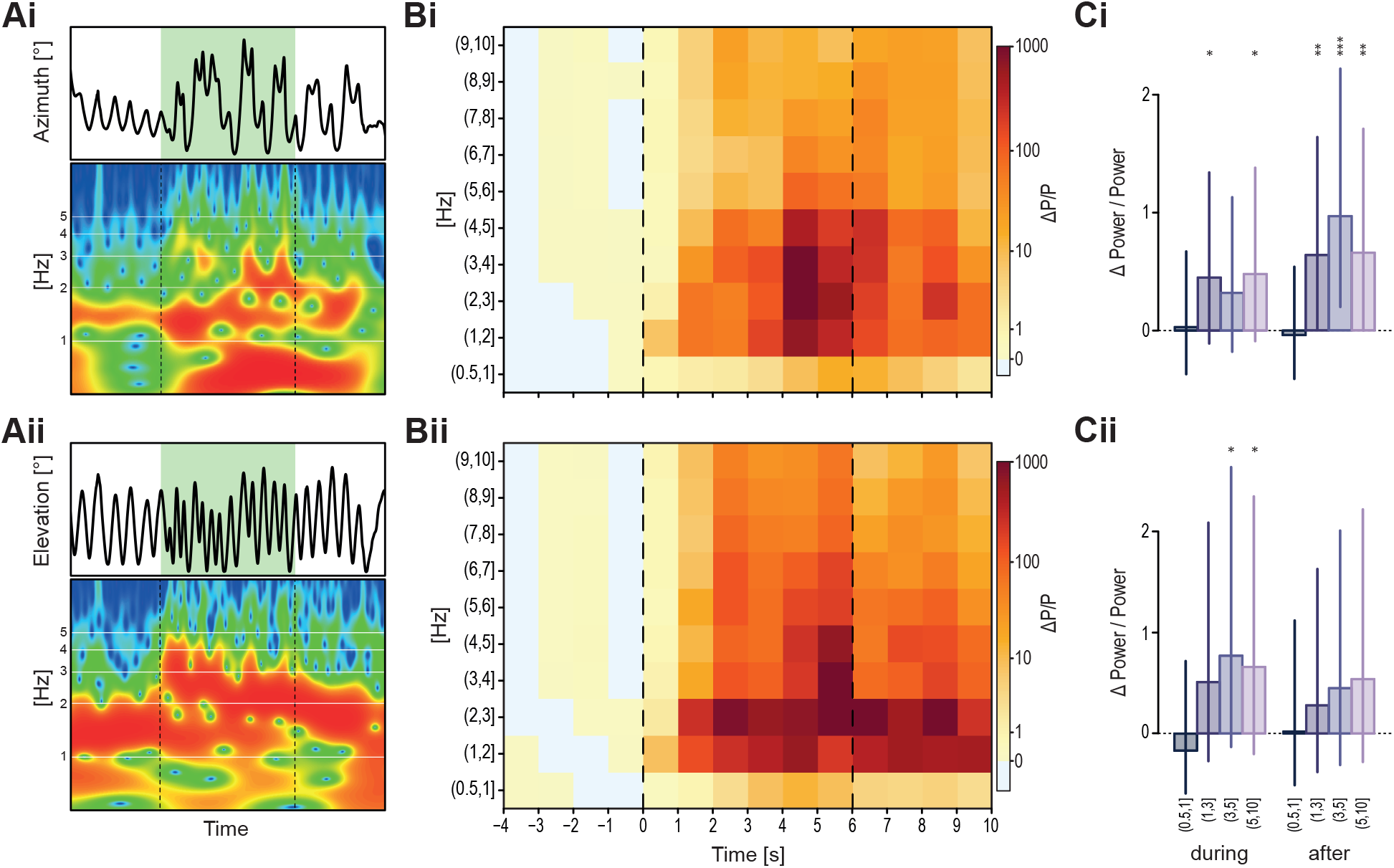
Odour-induced high-frequency antennal movements. (A) A single trial example with time series and its corresponding wavelet transform for the antennal angles in azimuth (i) and elevation (ii). Shown is a truncated log frequency range of 0.5 – 10 Hz. Green shading indicates the time of the odour stimulus. (B) Mean change in wavelet power, scaled for each frequency band by its pre-stimulus average of antennal angles in azimuth (Bi) and elevation (Bii). Vertical dashed lines indicate odour onset and offset (Col).(C) Corresponding model means and credible intervals of scaled power changes (ΔP/P), during and after the odour stimulus (Col). For the analysis, frequency bands were pooled in 4 frequency ranges, showing separately the analysis for the Horizontal (i) and the vertical (ii) movement components. N=24 for azimuth, N=14 for elevation. Asterisks indicate the certainty levels of each mean to be different from zero (*: ≥ 90%, **: ≥ 95%, ***: ≥ 99%).

For all metrics above, **odour-induced effects** were calculated by comparing metrics between the odour time window and a time window directly before the odour onset (of the same length). In experiments with a shifting stream (Fig. 4) we had a sham control with a non-odorised air stream. In addition to the pre-stimulus baseline, we corrected the metrics of each odour-stimulus trial with the values of the sham control trial of the same animal.

### 4.5 Responsive vs non-responsive antennae

For the central stream condition (Fig. 2 – position 1 - C, Di), we defined the “responsive antenna” as the one with the largest decrease in distance to the stream (regardless of its left/right identity). As a control, we applied the same method to a baseline variation between two consecutive time windows before odour onset (only clean air) and without transferring antenna identities (grey lines). This allowed us to compare the inter-antennal differences with a neutral baseline representing the natural asymmetries in movement. The identity of the antenna labelled as “responsive” in terms of distance to the stream was then kept for the antennal range calculation, which was then compared with its antenna-specific baseline variation instead of a neutral baseline. In the conditions with asymmetric stimuli (Fig. 3 – position 2 & Fig. 4), the left/right identities were maintained for all quantifications.

### 4.6 Visualising air-borne plumes

TiO_2_ smoke resulting from the contact of TiCl_4_ (Fluka, CAS 7550-45-0) with air humidity was used to visualize the air flow from the odour delivery device at a low wind speed (0.05 m/s) (Fig.S 4). A planar infrared laser (0.2mm light sheet thickness, 808nm wavelength, 7W, Laserwave, Beijing, China) was used to highlight a horizontal section of the smoke and reveal its flow patterns. Cockroaches were then placed in the wind tunnel, in an analogous manner to the odour experiments. A top-view of the laser-illuminated region was recorded with the top single Basler camera of the Motif system at 100 fps. Sequences of interest, where an antenna moved through the smoke (Fig. 6, Fig.S 4) were isolated for analysis. For the condition with a static antenna, a recently dead animal was used.

**Figure 6.**
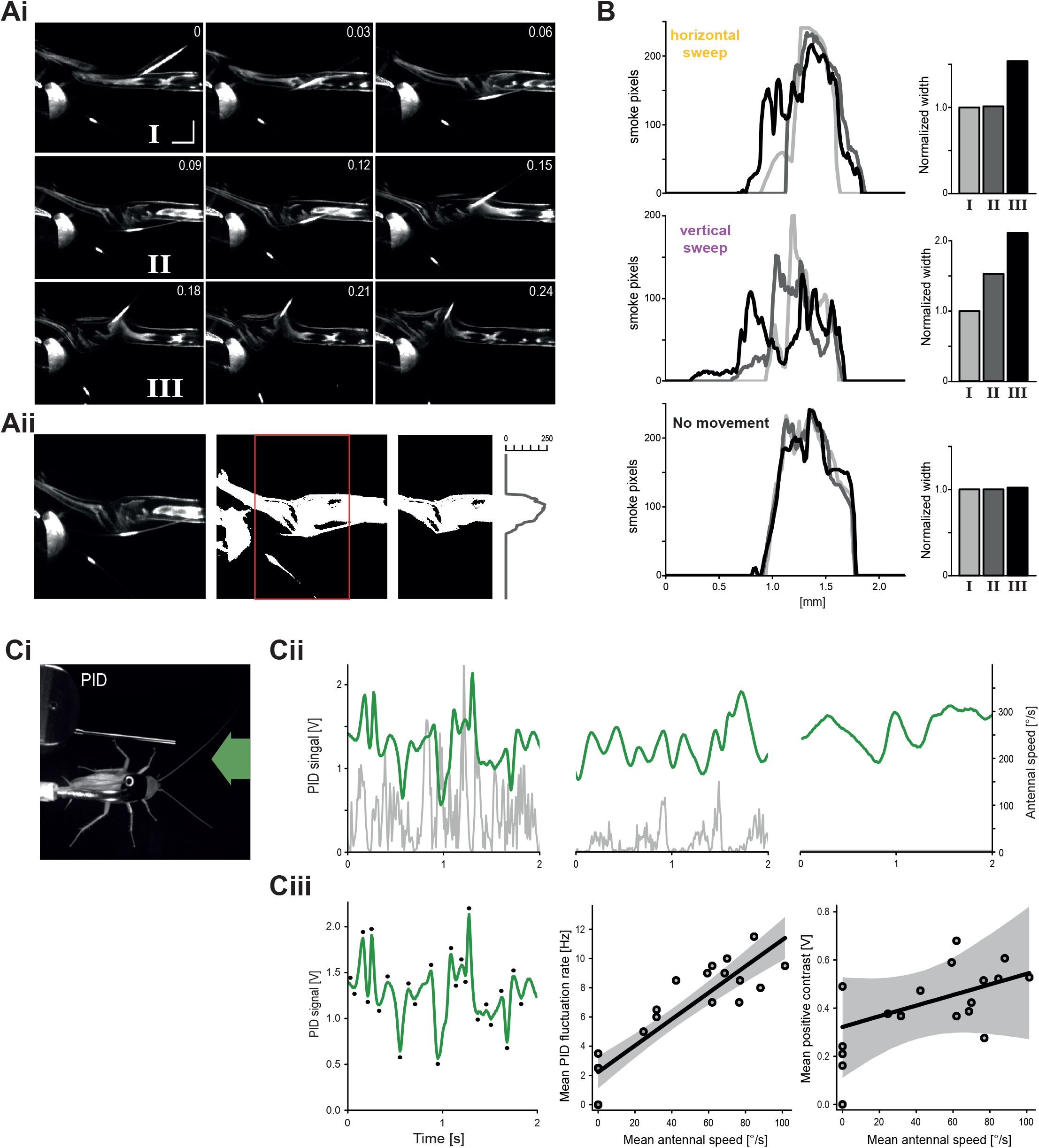
Antennal movement affect the distribution and dynamics of air-borne plumes. (A) Visualised air flow with TiO_2_ smoke. (i) Selected frames from a recorded sequence containing a horizontal antennal sweep through the plume. Time-stamps are in seconds, scale bars equal 10mm. The inward movement is seen between 0.03 – 0.09s, the outward movement between 0.12 – 0.21s. (ii) Illustration of the analysis process to estimate smoke distribution. From left to right: Raw frame, thresholded frame with the region of interest in front of the head marked in the red rectangle, removed antennae and limbs and the final distribution of smoke pixels within. (B) Smoke distributions for the three marked time-stamps I-III in (Ai) with their corresponding normalised widths, calculated according to Aii. Shades of grey represent time-stamps(I-III). Top: Frames during the horizontal sweep shown in (A). Frame I is before the sweep, frame II at the end of the inward motion, and frame III at the end of the outward sweep. Middle: A fast vertical sweep shown in Fig.S 4B. Here frame I is before the sweep, frame II at the end of a full sweep cycle and frame III is 0.09s afterwards. Bottom: A control sequence with a static antenna in the smoke plume (Fig.S 4C). (C) Simultaneous antennal movement tracking and PID recording as a proxy for odour fluctuations. (i) Recording setup with the position of the PID, animal and odour stimulus. (ii) Example recordings of the PID signal in green and simultaneous angular antennal speed in grey, for three different antennal movement regimes: fast, slow and a stationary control with a fixed antenna. (iii) Left: An example PID trace with automatic peak detection to estimate rate and amplitude of odour fluctuations. Middle: Correlation between the mean PID voltage peak rate and mean antennal speed during 2s time windows. Right: Correlation between the mean positive voltage contrasts (signal increase from a negative peak to the next positive peak) and the mean antennal speed for the same 2s windows. In both, solid lines show linear regression curves with their shaded credible intervals (n = 22).

#### Smoke distribution

Selected frames were processed in R and Adobe Photoshop. We carefully hand-drew a mask on the animal using the original grayscale image in order to have a silhouette. We thresholded the original grayscale images with the same threshold value for all frames of a given sequence and defined a region of interest (ROI) for analysis in front of the head. We then applied the mask to the thresholded images in order to remove pixels representing body parts and antennae. This process was done manually, resulting in cropped images where white pixels only represent smoke particles. We then calculated the number of white pixels for each row of the ROI (In R, https://cran.r-project.org/), giving us the left-to-right distribution of smoke particles in front of the animal (Fig. 6).

### 4.7 Odour fluctuations and antennal movements

As a proxy for the relative odorant concentration over time during antennal movements, the PID sensor was placed at head level, just behind the left antenna in its natural resting position (fig. 6B). The PID pump was set to its lowest setting and air speed to 0.05m/s. The odour stream was centered between the head and the sensor (at this location and air-speed, baseline fluctuations were more pronounced than in fig. 1). The odorant used was pure butanol (200μL in a 20mL vial), as it is easily detected by the PID (butanol ionisation potential = 10.04 eV). **Tracking** of the head and antenna tip was achieved with the loopy software (LoopBio GmbH, http://loopbio.com/loopy/), from a top view. Azimuth and elevation angles were used to compute the combined angular antennal speed in R.

#### Correlation between antennal movements and odour fluctuations

The analysis was performed using 2-second windows, selected systematically according to local peaks in antennal speed (after a moving average filter), resulting in a wide range of antennal movement regimes. As a control, windows of the same size were randomly selected from recordings with the antenna fixed at its base. Odour fluctuations were estimated from the rate and magnitudes of the PID voltage peaks (fig. 6B). All calculations for peak detection, fluctuation rate (average instantaneous frequency of all detected PID minima and maxima) and magnitude (average voltage difference between minima and their subsequent maxima - positive contrasts) were performed in R.

### 4.8 Statistical analysis

Throughout, we used Bayesian statistics based on Markov Chain Monte-Carlo sampling (using the *brms* (Bürkner (2017)) and the *cmdstanr* (Gabry and Češnovar (2022)) packages, implementing *Stan* in R (Stan Development Team (2022)) in combination with methods from Korner-Nievergelt et al. (2015)). We used weakly informative priors from the *brm* function. All model outputs are presented in the scale of the original data and were thus back-transformed if any transformation (pre-transformation and/or link transformation) was used to fit the model. Estimates correspond to 50% quantile of the posterior distributions (mean), with the 2.5% and 97.5% quantiles as 95% credible intervals. Certainty of odour-induced effects and comparisons were tested by calculating the proportion of posterior samples that were above or below a reference value of either zero (i.e. the pre-stimulus baseline), a control value, or another within-model group. Differences with ≥90%, ≥95%, and ≥99% certainty are marked with asterisks (see figure legends for details on the comparisons made in each). Table 1 summarises the model parameters for each analysis.

**Table 1.**
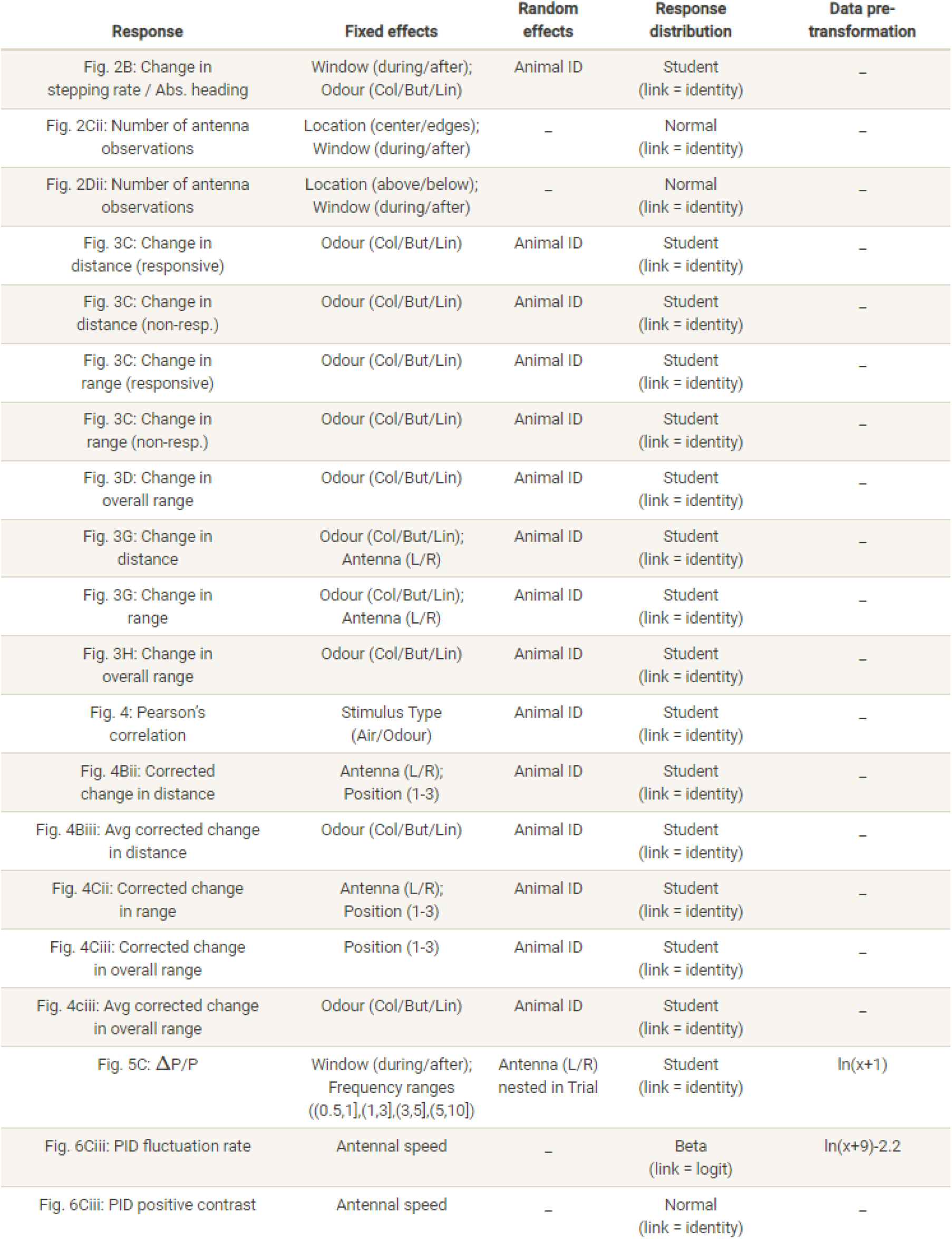

### 4.9 Data and code availability

Data was analysed and plotted using custom-written R scripts. Analysis code and data is available at https://github.com/Couzin-Fuchs-Lab/ActiveSmellingCockroach.

## 5 Results

### 5.1 Movement kinematics with odour exposure

Antennal kinematics were measured under laminar flow conditions in our wind tunnel (Fig. 1Ai) in order to characterise whether and how movement dynamics are changed according to the olfactory environment. Overall, we performed 215 trials on 31 animals, tracking their legs, head and antennal tip positions in 3D (Fig. 1B) to characterise their kinematics. The general analysis, pooling trials and conditions, revealed continuous horizontal antennal sweeping that includes relatively large and slow cycles (Fig. 1C) and a characteristic double tripod walking gait (Fig. 1D). Animals were tested with three test odours delivered in different positions. The set of odours, comprising of an innately attractive colony extract mixture, a mildly-appetitive short-chain alcohol butanol and a neutral to mildly aversive single-molecule floral odorant linalool (see Fig.S 1B for a preference test), were chosen as a relatively broad set of odours to highlight both common features and differences in observed responses.

Shortly after odour presentations, changes in legs and antennae kinematics were observed. A single trial example of antennal movements (horizontal and vertical components of the left antennal tip) and the corresponding stepping rate and heading direction during a 6 seconds odour stimulus (colony, central position 1) are shown in (Fig. 2Ai). As seen here and in most tested animals, shortly after odour onset, animals slowed down leg movements and centrally aligned their heading direction. Fig. 2B shows time traces and binned averages for trials with the colony odour. Responses for the other tested odors, in Fig.S 2C and D, show a similar decrease in leg movements but no consistent immediate change in heading. In parallel, changes in antennal position were observed in both azimuth and elevation (Fig. 2C and D). In the absence of odour, the antennae sweep around their natural position of approximately 50^*?*^ from the midline, covering a wide range with a relatively small overlap at the centre (left heatmap in Fig. 2Ci and grey density curve in Fig. 2Cii). When an odour is presented around the midline position (Fig. 2A, stimulus in position 1), this overlap increases (Fig. 2Ci right heatmap and green density curve in Fig. 2Cii). In addition, the antennal sweeps, which are mostly confined to ground level in the absence of odour, show drastic increases in elevation, covering a larger vertical space with more frequent upstroke movements (Fig. 2D). This behaviour is not odour-specific, with similar changes in coverage observed for the other tested odours (details in the subsequent figures below and Fig.S 1E-F).

### 5.2 Trade-off between local sampling and large-scale scanning

Interestingly, even though both antennae had equal access to the odour, the increase in activity around the odourised stream predominantly occurred in one of the two antennae (Fig. 3B, low-level of pre-stimulus asymmetry (dashed-line curves) is increased during the odour presentation (solid line curves)), with no consistent side bias across trials (Fig.S 2B). Note that, as the stream is centred, the proximal end of both antennae were in contact with the odour throughout the trials. In addition, in the majority of cases, the same ‘responsive’ antenna also increased its sweeping range (Fig. 3C) which resulted in longer time spent around the odour without compromising the overall sensory space (Fig. 3D shows the overall space covered by both antennae in unaffected). To test whether this behaviour is specific to the central location, a similar analysis was carried out with a second set of trials in which the odour stream was positioned on the right side of the tunnel (Fig. 3E-H, position 2). In this position, the contralateral antenna (in red) cannot reach the stream. As for position 1, when an odour was presented, the antennae spent more time overall close to the odour (Fig. 3E-G). Here, the contra-lateral antenna increased its sweeping range, thus reaching further towards the odour side, while the ipsilateral one (in blue) tended to decrease its range around the odour (Fig. 3G). Altogether, however, as for the centre plume, the overall scanning range covered by both antennae was mostly maintained, for all three tested odorants (with a slight decrease in range for the side stream with linalool, Fig. 3H). This represents an effective space coverage, in which, via asymmetric scanning behaviour (spontaneous in position 1 and experimentally imposed in position 2), an increased odour exposure is achieved without a loss in exploration range for both stream positions.

### 5.3 The antennae follow a moving odour stream

To further test these findings, we monitored antennal movement during moving stimuli (Fig. 4), starting from the centre position 1, via the intermediate position 2 (side stimulus in Fig. 3) to the very edge of the animal’s reach (dashed white/green area in Fig. 4A shows the air/odour positions for this experimental set, noted also as P1-P3). To differentiate between mechanical and odour-induced effects, all responses of odour trials were compared to sham air controls. Analysing body movement reveals significant odour following behaviour in heading direction (bottom panel in Fig. 4A shows that the average heading direction follows the plume movement in colony trials), and in walking behaviour (the initial decrease in speed with odour presentation in P1 changes to increased walking when the odour is shifted, middle panel Fig. 4A). This following behaviour was odour-specific, with clear following of colony and butanol, no following with air-control and linalool (Fig.S 2). This was reflected by the antennal activity, in which clear following behaviour in both antennae was observed for the attractive odours (significantly higher correlation between the movement of the antennae and the stream in presence of an odour compared with air control, see example in Fig. 4A and Fig.S 2C for correlation coefficient values). As illustrated in the example, the time course averages (Fig. 4Bi) and binned averages (Fig. 4Bii), with odour presentation, both antennae shifted their centre of sweep along with the stream’s movement such that a small distance to the odour was maintained throughout (Fig. 3Bii for P1-P3). Note that the antennal shifts are most salient at the positions in which they fully loose contact with the odour (P2 for the contralateral antenna and P3 for the ipsilateral one), potentially as a means to regain contact at the tip. As was the case for static streams, the sweeping range of the contralateral antenna (red) increased while the range of the ipsilateral one tended to decrease (Fig. 3G), with the overall range covered by both remaining unaffected (Fig. 3H and Fig. 4Ciii).

### 5.4 High-frequency antennal sweeps during odour encounter

To further characterise antennal spatio-temporal dynamics during odour encounters, we applied a wavelet analysis, decomposing the time-series of azimuth and elevation angular antennal positions into time-frequency space (example of horizontal and vertical movement traces with their respective wavelet decompositions are plotted in Fig. 5Ai and Fig. 5Aii, See Methods for details). This allows quantification of changes in sweeping patterns at multiple temporal scales simultaneously. As illustrated in the example trace (Fig. 5A) and population average (Fig. 5B and C), odour presentation is associated with an increase in power in all frequency bands above 1 Hz, both in azimuth and elevation for all tested odours (see also Fig.S 3). The increase in high frequency movement bouts lasts for the entire duration of the odour stimuli. Notably, the increase in horizontal sweeping persists after offset (and even further increases for the colony odour), while the increase in vertical movement is most significant during the odour stimulus (Fig. 5Ci and Cii).

### 5.5 High-frequency antennal movement impacts the odour distribution

Finally, we address the impact of antennal movement on the odour environment. We started with a simple quantification of the plume structure in different movement regimes. This was done by simulating an odorant-carrying air flow at low speed with light smoke. A planar infrared laser was used to visualise a horizontal section of the stream at the level of the animal, thus highlighting the effect of antennal movements on the air flow (see Methods and example snapshots in Fig. 6A and Fig.S 4B-C). We find that both slow horizontal and fast vertical sweeps affect the smoke distribution (top and middle panels in Fig. 6B show the distribution of smoke around the antenna - before, during and after a horizontal and vertical sweep respectively). The antennae, by successively passing back-and-forth through the stream, leave behind gaps when capturing smoke particles in their wake. The resulting distribution of the smoke after even a single sweep cycle is altered (Fig. 6A and B), an effect that lasts beyond the end of the sweep (Fig.S 4B). To further evaluate the movement impact on odour dynamics, a photo-ionisation detector (PID) was placed in the plume (Fig. 6Ci), close to the proximal third of the antenna, where the density of olfactory receptors is high (Paoli et al. (2020)). Continuous recording of PID voltage traces, a proxy for odour concentration (Fig. 6Cii, green traces), with the corresponding antennal kinematics (Fig. 6Cii, grey traces) reveal that odour fluctuations increase with increasing antennal movements (Fig. 6Ciii). Peak-detection analysis of the PID traces (marked by black dots in Fig. 6Ciii) demonstrates that concentration fluctuations increase in rate (mean instantaneous rate of peaks) and amplitude with increasing antennal speed (Fig. 6Ciii). Taken together, these findings highlight that antennal movements not only impact the rate of odour encounter by changing their position and sweeping pattern within an odour environment, but also by altering its structure.

## 6 Discussion

In this study, we investigated the use of antennal movements in olfactory sensing. We have shown that cockroaches track static and moving odours by shifting the antennae towards the plume center where local high-frequency movement bouts are carried out. Interestingly, however, long-range scanning sweeps continue throughout, such that the overall sensory range covered is not significantly impacted. Upon odour detection, a general increase in fast movement components was observed, with rapid vertical strokes exhibited most prominently during the stimulus duration, and increased horizontal sweeps that also continued after stimulus termination. A similar increase in oscillation frequency has recently been reported in bumblebees (Claverie et al. (2021)), corresponding to frequencies of optimal odour capture rates (Claverie et al. (2022)). Utilizing both numerical simulations and Particle Image Velocimetry (PIV) experiments with artificial antennae, Claverie et al. (2022) (see also Su et al. (2019) and Reidenbach et al. (2008)) demonstrated that the air flow vortexes generated by antennal vibrations can enhance the transport and adsorption of odour molecules through the sensory pores. In agreement with that, our *in-vivo* visualisations of the flow and odour concentrations around the cockroach antennae provide empirical evidence that even these thin appendages create significant flow fluctuations.

### 6.1 Exploration-exploitation trade-off

Our first aim was to investigate how antennal movement participates in spatial sampling. It has recently been shown that honeybees orient their antennae towards appetitive odour sources and away from aversive ones (Cholé et al. (2022); Gascue et al. (2022)). Moreover, the analysis demonstrated a general trend of positive correlation between the time the antennae spent focused on an odour target and its valence (Gascue et al. (2022)). Similarly, we also observed that cockroaches shift the sweeps of at least one antenna towards the stream of appetitive odorants (exemplified by a decrease in distance to the odour stream, regardless of the stream’s physical location). In terms of optimising the information animals can extract from their environment, there is an interesting interplay between sensing, or localising a particular source of interest (‘exploitation’), and scanning the rest of the sensory space (‘exploration’). Our observations suggest that for antennal movement, this trade-off can be resolved with asymmetric sweeping, such that one antenna in particular spends more time in the odourised region. For central stimuli where there is no inherent cause for asymmetry, there was no consistent left/right bias across trials and animals (ref SI). Regardless of the stimulus position, for both stationary and moving odours, the sweeping range of one antenna is adjusted such that the overall scanning range (and therefore the possibility to detect other stimuli elsewhere) is maintained. This behavioural asymmetry, manifested differently depending on whether the stimulus is central or not, may offer a natural ‘division of labour’ configuration for efficient local sensing and larger-range scanning.

### 6.2 Odour intermittency and parallels to sniffing

Temporal dynamics of odour encounters depend on both the spatio-temporal structure of the odour environment as well as the way an organism moves, interacts and potentially reformats its olfactory surrounding (Crimaldi et al. (2021); Lei et al. (2022)). Intermittency and fluctuations in odour plumes have been shown to tune the neural activity in the peripheral olfactory circuits (Huston et al. (2015); Lei et al. (2009)) and were suggested to improve up-wind odour-source localisation (Lei et al. (2009); WILLIS and BAKER (1984)). In addition to natural fluctuations, animals often display small oscillatory movements on top of their primary task-related trajectories (Chen et al. (2020); Liberzon et al. (2018); Willis and Avondet (2005)), which are intensified with novel stimuli detection (Huston et al. (2015); Loudon and Koehl (2000); Schmitt and Ache (1979)). These behaviours enhance odour reception by imposing frequency-dependent fluctuations and creating sharp odour onset at the olfactory organs (Antennule ‘flicking’ in lobsters, Koehl et al. (2001); wing-flapping in flying insects, Li et al. (2018); Loudon and Koehl (2000) and antennal movement, Claverie et al. (2022); Huston et al. (2015)). We show here that cockroaches display comparable odour-induced modulations of antennal movement (Fig. 5, Fig.S 3) impacting the spatio-temporal structure of the odour environment (Fig. 6, Fig.S 4), which could be of particular importance in low-turbulence environments where cockroaches dwell. These elements can be paralleled with mammalian sniffing (Verhagen et al. (2007)). During sniffing, the sharp input events, created through the rapid flow of odorants into the nasal cavity, have been shown to enhance odorant detection and discrimination (Kepecs et al. (2007); Oka et al. (2009)). The sniffing frequency is controlled on a cycle to cycle basis according to task, the complexity of the olfactory environment and the animals’ internal state (Verhagen et al. (2007)) and increases upon odour encounter (Bhattacharjee et al. (2019); Reisert et al. (2020)). At low to intermediate sniffing rates, intermittency in odour concentration allows receptor neurons to recover from adaptation during the intervals between successive inhalations Wachowiak (2011). With increase in sniffing frequency, seen during active exploration, signal attenuation also increases, resulting in enhanced contrasts for target odorants (temporally dynamic or spatially localized) against the broadly distributed background odours (Verhagen et al. (2007); Wachowiak (2011). Parallels in the functional neural architecture of different olfactory systems (Strausfeld and Hildebrand (1999); Touhara and Vosshall (2009)) further suggests that movement-driven temporal patterning in arthropods could similarly impact odour coding in a frequency-dependent manner.

## 7 Acknowledgements

The authors would like to thank Liang Li for help with setting up the infrared laser system and Jahn Nitschke, Alejandro Pequeno-Zurro, Paul Szyszka and Loopbio GmbH for their contributions in setting up the wind tunnel experiments. Thanks also to Jaclyn McCollum and Hannes Kübler for contributing to the data collection on odour preference (Supplementary Figure 1A).

This work was completed with the support of the Deutsche Forschungsgemeinschaft (DFG, German Research Foundation) under Germany’s Excellence Strategy – EXC 2117 – 422037984.

## 10 Supplementary Information

**Figure 1.**
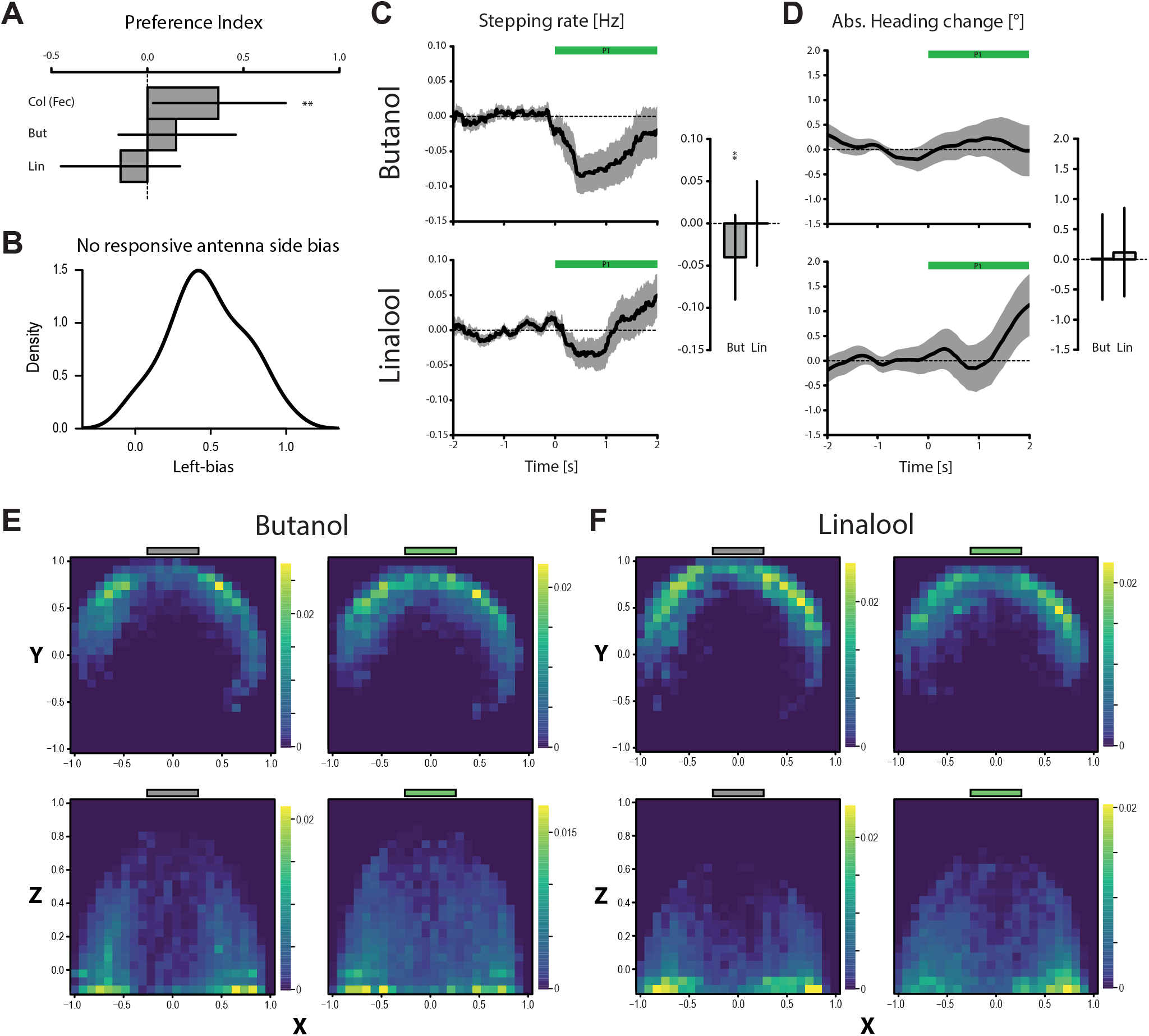
Supplementary figure for centre stimuli. (A) Preference index (PI) computed from arena choice-assays between an odourised shelter and a control shelter: *PI* = (*Time*_*odour*_ *Time*_*ctrl*_)*/Time*_*total*_ (N=15 for Colony(Fec), N=19 for Butanol and Linalool. As a colony odour, a cockroach feces extract was used (provided by Hiroshi Nishino, Hokkaido University, Japan). See Günzel et al. (2021) and the provided analysis code for more details. (B) Distribution of left-biases for the identity of the responsive antenna across trials for all odours, within animals: *Lbias*_*animalID*_ = *N*_*Lresp*_*/*(*N*_*Lresp*_ + *N*_*Rresp*_) (n=138 trials, N=26 animals). (C) Change in stepping rate (n=38 trials, N=27 animals for Butanol; n=45 trials, N=24 animals for Linalool). (D) Change in absolute heading (n=46 trials, N=30 animals for Butanol; n=48 trials, N=28 animals for Linalool). Mean time-series ± s.e.m. and model means with credible intervals. In (A)-(C), asterisks indicate the certainty levels of the means to be different from zero (*: ≥ 90%, **: ≥ 95%, ***: ≥ 99%) (E) Top row: Horizontal antennal tip density maps in the normalised coordinate system (zero = head position, ±1 = maximum coordinate across trials) before (left, grey bar) and during (right, green bar) a butanol stimulus (n=46 trials). Colours indicate relative proportions of observations per pixel. Bottom row: Same thing for the vertical antennal tip presence density in the normalised coordinate system (zero = head position, 1 = maximum coordinate across trials, 0 = ground). (F) Same as in (E) for a linalool stimulus (n=48 trials).

**Figure 2.**
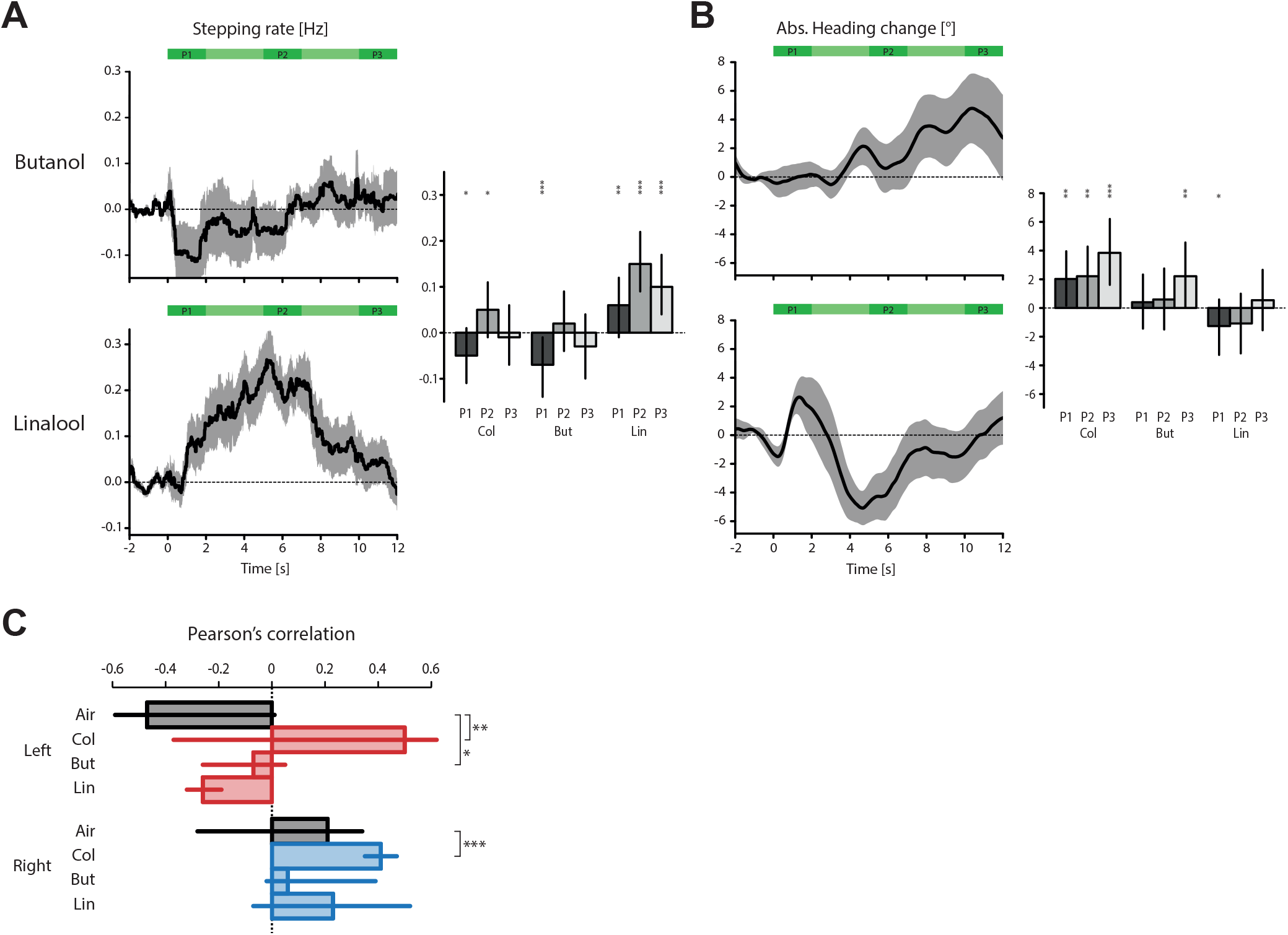
Supplementary figure for moving stimuli. (A) Change in stepping rate (n=13 trials for Colony; n=10 for Butanol; n=12 for Linalool). (B) Change in absolute heading (n=13 trials for each). Mean time-series ± s.e.m. and model means with credible intervals for each odour and each static position (P1-3). Asterisks indicate the certainty levels of the means to be different from zero (*: ≥ 90%, **: ≥ 95%, ***: ≥ 99%). (C) Pearson’s correlation coefficients between the antennal position and the stream centre for each antenna and each odour, as well as the air sham control (N=13 trials for each). Asterisks indicate the certainty levels of the antennal means to be different from their respective controls (*: ≥ 90%, **: ≥ 95%, ***: ≥ 99%).

**Figure 3.**
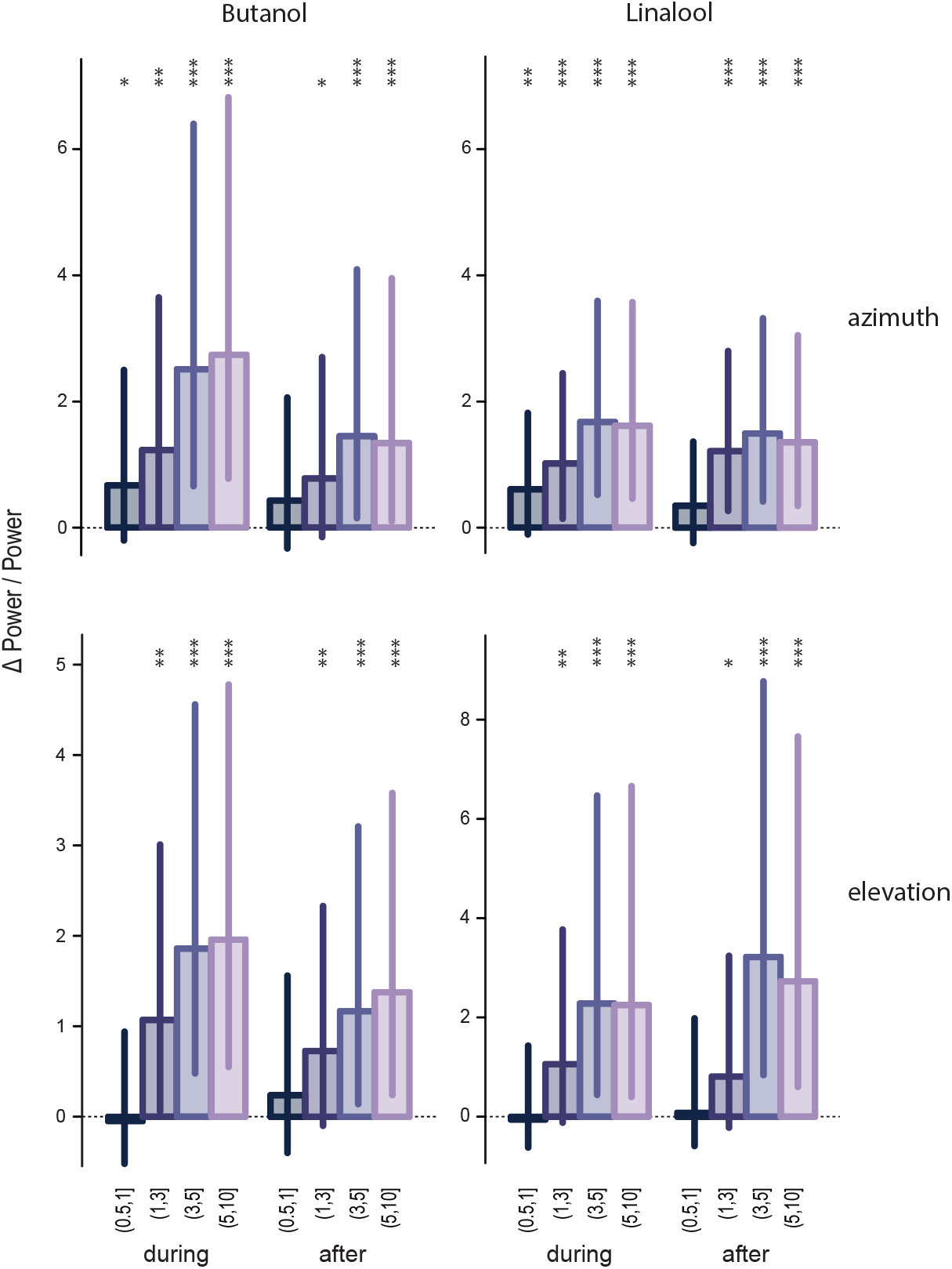
Supplementary figure for the wavelet analysis. Model means and credible intervals of scaled power changes (ΔP/P), during and after the odour stimuli (left: butanol, right: linalool). Frequency bands were pooled in 4 frequency ranges, showing separately the analysis for the Horizontal (top row) and the vertical (bottom row) movement components (N=13 for butanol; N=26 for linalool azimuth, N=14 for linalool elevation). Asterisks indicate the certainty levels of each mean to be different from zero (*: ≥ 90%, **: ≥ 95%, ***: ≥ 99%.

**Figure 4.**
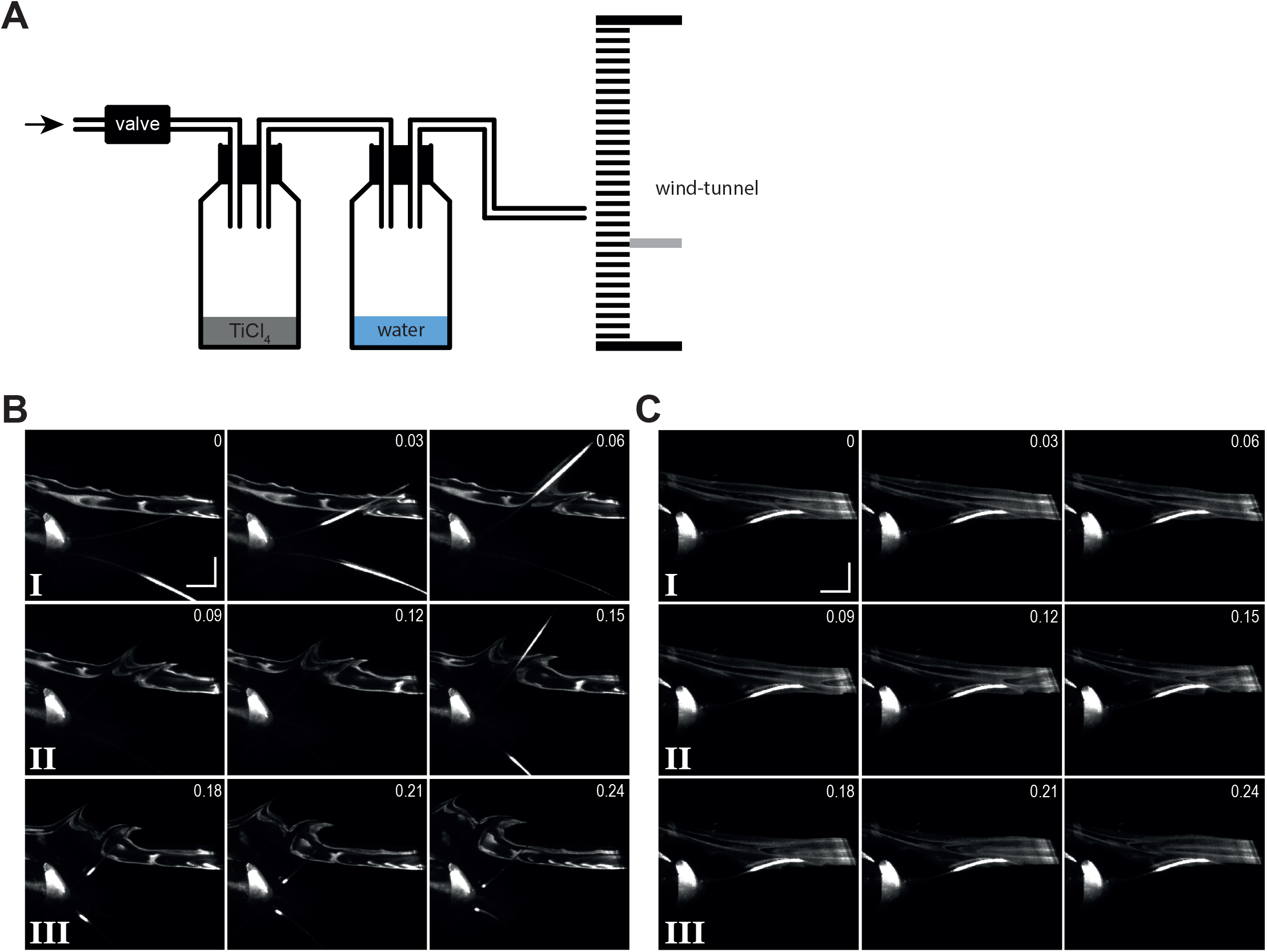
Supplementary figure for the plume visualisations. (A) Schematic of the smoke delivery set-up. (B) Visualised air flow with TiO_2_ smoke. Selected frames from a recorded sequence containing a vertical antennal sweep through the plume. Time-stamps are in seconds, scale bars equal 10mm. The upward movement is seen at 0.03 and 0.06s, the downward movement is already completed at 0.09s. Frames labelled I-III are analysed in Fig. 5B. (C) Same thing with a dead animal for the ‘no movement’ control.

